# Improving rigor and reproducibility in chromatin immunoprecipitation assay data analysis workflows with Rocketchip

**DOI:** 10.1101/2024.07.10.602975

**Authors:** Viktoria Haghani, Aditi Goyal, Alan Zhang, Osman Sharifi, Natasha Mariano, Dag Yasui, Ian Korf, Janine LaSalle

**Affiliations:** Department of Medical Microbiology and Immunology, Genome Center, University of California, Davis. Davis, CA, USA; Department of Molecular and Cellular Biology, Genome Center, University of California, Davis. Davis, CA, USA

**Keywords:** ChIP-seq, CUT&RUN, CUT&Tag, bioinformatics, workflow, Snakemake

## Abstract

As genome sequencing technologies advance, the accumulation of sequencing data in public databases necessitates more robust and adaptable data analysis workflows. Here, we present Rocketchip, which aims to offer a solution to this problem by allowing researchers to easily compare and swap out different components of ChIP-seq, CUT&RUN, and CUT&Tag data analysis, thereby facilitating the identification of reliable analysis methodologies. Rocketchip enables researchers to efficiently process large datasets while ensuring reproducibility and allowing for the reanalysis of existing data. By supporting comparative analyses across different datasets and methodologies, Rocketchip contributes to the rigor and reproducibility of scientific findings. Furthermore, Rocketchip serves as a platform for benchmarking algorithms, allowing researchers to identify the most accurate and efficient analytical approaches to be applied to their data. In emphasizing reproducibility and adaptability, Rocketchip represents a significant step towards fostering robust scientific research practices.

## Introduction

As genome sequencing technologies and their applications continue to rapidly evolve to include epigenomic information, a vast amount of sequencing data is accumulating (1). Journals now commonly mandate the deposition of raw sequence data to public databases, such as the International Nucleotide Sequence Database Collaboration Sequence Read Archive (SRA), generating a substantial volume of sequencing data (2). These mandates are further supported by funding agencies, such as the National Institutes of Health (NIH), which requires NIH-funded research to publish sequence data as part of their Genomic Data Sharing Policy (3). Therefore, there is an increasing need for more comprehensive and biologically relevant data analysis workflows that promote reproducibility of results and allow for increased leverage and comparisons of publicly available data. Although mandated availability of data is beneficial to science, data analysis pipelines are often complicated, with divergent results due to variation in analysis steps, parameter usage, software used, and software version. These problems can partly be solved by workflow managers, which control for analysis step order, software versions via virtual environments, software parameters, etc. This is especially important for workflows requiring multiple analysis steps, such as sequence data produced by chromatin immunoprecipitation assays, where small differences in analytical steps can yield different results.

Chromatin immunoprecipitation followed by sequencing (ChIP-seq) is a technique commonly used to identify protein binding sites in the genome (4,5). Briefly, DNA associated proteins are chemically fixed onto DNA via a crosslinking agent. The resulting chromatin is then fragmented by either sonication or enzymatic digestion into 100-500 base pair fragments. Next, chromatin fragments bearing the protein of interest are immunoprecipitated using protein-specific antibodies. Typically, ChIP-seq experiments utilize two controls: (1) an input control to correct for differences in sonication and genomic DNA sequence bias and (2) a mock IP to account for nonspecific interactions of the antibody used (6–8). Finally, the chemical cross-link is reversed, allowing the DNA bound by the protein to be purified and then sequenced. In addition to ChIP-seq, other chromatin immunoprecipitation assays, such as cleavage under targets and release using nuclease (CUT&RUN) and cleavage under targets and tagmentation (CUT&Tag) are similarly utilized to assess protein binding sites (9,10). In CUT&RUN, antibody binding occurs directly on protein-bound DNA fragments within intact nuclei, with DNA fragmentation accomplished enzymatically using a fusion protein containing Protein A and micrococcal nuclease (MNase), contrasting with ChIP-seq’s reliance on sonication for DNA fragmentation. In CUT&Tag, a fusion protein combines Protein A and Tn5 transposase, allowing for simultaneous tagging and binding of sequencing adapters to protein-bound DNA fragments.

Following the unique approaches of ChIP-seq, CUT&RUN, and CUT&Tag in isolating protein-bound DNA, the generated DNA sequence data is aligned to a reference genome, and a peak caller is typically used to identify regions of interest. These peaks are “broad” for proteins with large areas of interactions with DNA, such as histones and DNA methylation interacting proteins. Conversely, proteins like transcription factors such as CTCF, that have a sequence specific region of interaction, produce “narrow” peaks. Analyzing these different types of DNA binding proteins can sometimes call for different computational controls. Given the variation in how peaks are generated, there is a significant amount of statistical noise in these experiments, which can hinder the efficacy of peak calling algorithms. This is further complicated by sequencing issues associated with strand bias, GC content, PCR amplification, library preparation, primer choice, sequencing platform, and antibody choice (8,11–20). Consequently, it is increasingly difficult to reproduce or replicate experimental findings. From this point forward, “reproducibility” refers to an individual’s or group’s ability to generate the same findings as another study, whereas “replicability” represents an individual’s or group’s ability to recreate findings multiple times using the same data input. Overall, this highlights the need for consistent and controlled data analysis practices, ensuring reproducibility and replicability of results.

Here, we present Rocketchip, available at https://github.com/vhaghani26/rocketchip, to address key aspects of these problems. Rocketchip reduces variation in data analysis methodologies, increase reproducibility and replicability of experimental results, and encourage greater usage of publicly available sequence data.

## Methods and Results

### Implementation

Rocketchip is an automated bioinformatics workflow written in the Python-based (v3.10.12) workflow manager, Snakemake (v7.32.4) (21) (Figure 1). Rocketchip downloads ChIP-seq data directly from the SRA, the largest publicly available sequence database, using the SRA Toolkit (sra-tools, 3.0.9) (2,22). The SRA Toolkit downloads data via the prefetch and fasterq-dump functions, then splits the raw DNA sequence read file into respective paired-end (PE) read files using the flag --split-files and converts them into FASTQ file formats, whereas for single-end (SE) read files, the SRA Toolkit is utilized solely for the download and conversion of raw read data into FASTQ format. Rocketchip also presents the option for users to use local (i.e. non-SRA sourced) ChIP-seq data by providing their own FASTQ files. In parallel, Rocketchip downloads and processes a reference genome of the user’s choice from the UCSC Genome Browser (23). After downloading and converting files, files are stored in the local file system in FASTA and FASTQ formats. Users also have the option of storing their own custom genomes and sequencing reads that are not yet publicly available.

**Figure 1.**
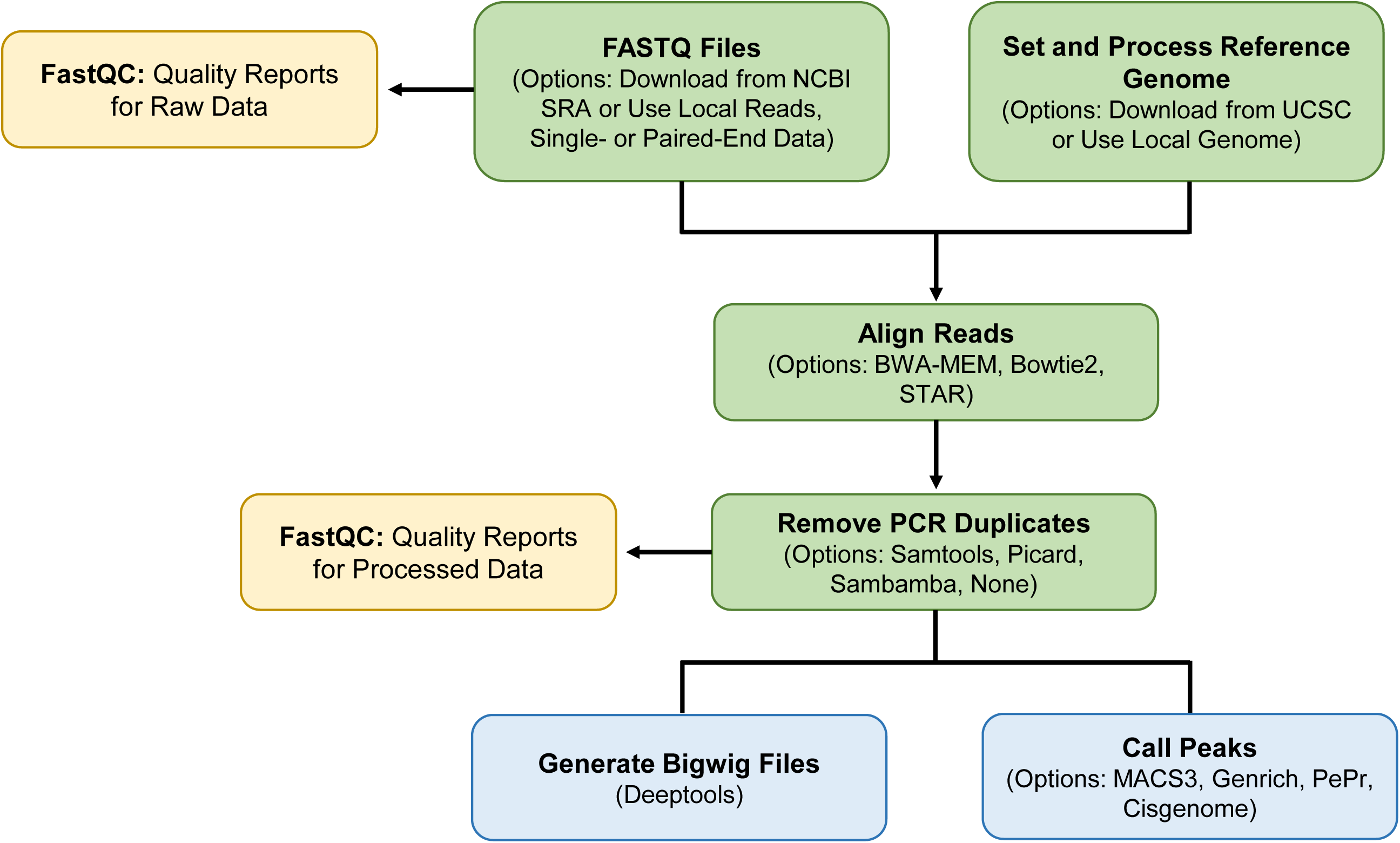
Rocketchip Pipeline for ChIP-seq Data Analysis. Raw sequencing data is aligned to a reference genome and processed to generate bigwig files for data visualization and delineate ChIP-seq peaks.

In Rocketchip, raw sequence data undergoes a quality control step using FastQC (v0.12.1) with default parameters to assess levels of data duplication and sequence quality (24). Raw sequence data is then aligned to the reference genome using the user’s choice of alignment software from BWA-MEM (v0.7.17), Bowtie2 (v2.5.2), or STAR (v2.7.11a), each with the default parameters (25–27). Intermediate data files are processed as necessary for deduplication. PCR duplicates are removed from sequence data using a user’s choice of deduplication software from Samtools (v1.18) with --mode s (i.e. standard PCR duplicate detection), Picard (v3.1.1) using MarkDuplicates, Sambamba (v1.0.0) with default parameters, or no deduplication (28–30). Deduplicated data files are subject to another quality control step using FastQC to ensure data integrity. Next, Deeptools’ (v3.5.4) bamCoverage function with default parameters is used to convert data from the BAM file format to the bigwig file format, which can be used for visualization of ChIP-seq data in the UCSC Genome Browser or other visualization tools (31). Finally, in Rocketchip, peaks are called with a user’s choice of software from MACS3 (v3.0.0b3) with the --bdg option, Genrich (v0.6.1) using default parameters, PePr (v1.1.24) using default parameters, or CisGenome (v2.0) (32–35). For CisGenome, if a control was used, the seqpeak command was employed with the default options. Without a control, CisGenome ran two rounds of peak calling using the hts_peakdetectorv2 command with options “-w 100 -s 25 -c 10 -br 1 -brl 30 -ssf 1.”

Peak-calling options in Rocketchip include narrow versus broad peak calling and use of a control where appropriate. Software version control is handled using Conda to ensure reproducibility of results (36). Software options were chosen based on options for command line use, ease of installation, and their standard use in the field.

### Validation of all Software Options

In order to ensure that all combinations of software can be integrated seamlessly, we selected a deeply sequenced experimental ChIP-seq study targeting a transcription factor, MeCP2, in mouse main olfactory epithelium conducted by Rube et al. 2016, hereafter referred to as “Rube”, (37) and a ChIP-seq study targeting NRF2 in human non-small lung cells conducted by Namani et al. 2019, hereafter referred to as “Namani” (38). To facilitate this comprehensive assessment, these data sets were run through Rocketchip using all four peak callers, namely MACS3, CisGenome, Genrich, and PePr. This was used in combination with each of the three aligners, BWA-MEM, Bowtie2, and STAR and four deduplication techniques, Samtools, Sambamba, Picard, and no deduplication. Additionally, to validate that Rocketchip can be run with or without a control (i.e. input or IgG control), this analysis was conducted with both the usage and omission of the corresponding control for each data set, with the exceptions of CisGenome and PePr, which can only be run if a control is used in the analysis. Each test was run three times to assess Rocketchip’s ability to replicate experimental results.

Each algorithm demonstrated varying peak-calling efficiency, influenced by several key factors (Figure 2). Most notably, the source of the data (i.e, Namani vs. Rube) revealed dramatic differences in the performance of peak-calling algorithms. For the Rube data, MACS3 consistently yielded the highest number of called peaks. All methods of deduplication produced comparable peak counts. Bowtie2 and BWA-MEM performed similarly, identifying slightly more peaks compared to STAR. CisGenome called the fewest peaks, significantly influenced by the deduplicator and aligner used. Without deduplication, CisGenome identified the fewest peaks across any software combination, while peak counts increased with any deduplication method. When STAR was used as the aligner with CisGenome, peak counts were the lowest. BWA-MEM yielded a higher peak count than Bowtie2 but exhibited more non-deterministic behavior compared to Bowtie2 and STAR. For Genrich, peak counts increased when the control was omitted during peak calling. In contrast to CisGenome, Genrich had the highest peak counts when no deduplication was used and when STAR was used for alignment. The other deduplicators and aligners showed negligible differences in peak counts. When using PePr for peak calling on the Rube data, no deduplication yielded the highest peak count, with other deduplication methods yielding similar results. Unlike the other algorithms, each aligner performed differently with PePr, with STAR identifying the most peaks, followed by BWA-MEM and then Bowtie2. Both the narrow- and broad-peak-calling algorithms yielded negligible differences, except for CisGenome, which had less deterministic results when peaks were defined as narrow and run through Rocketchip.

**Figure 2.**
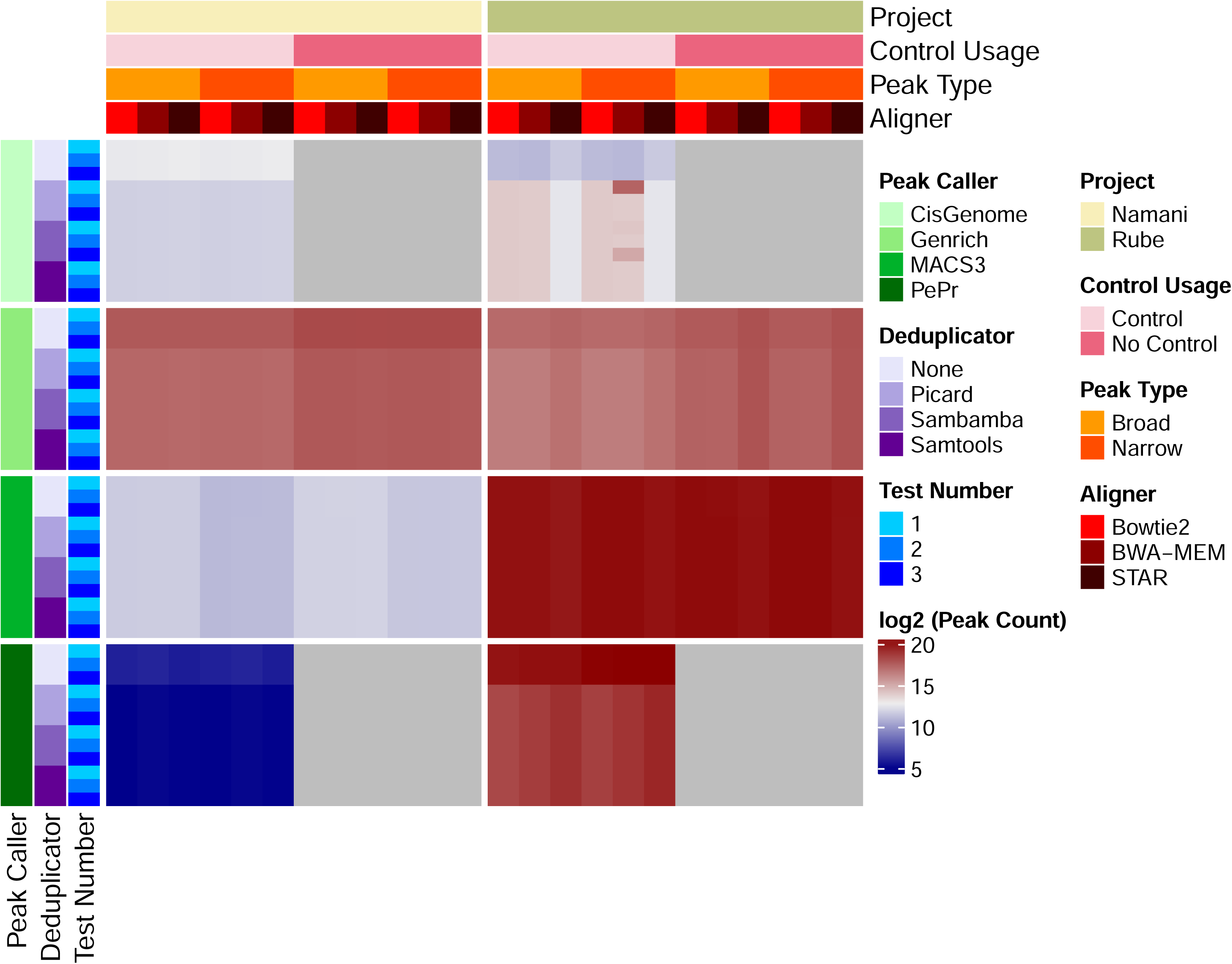
Peak Counts for all Software Combinations. ChIP-seq data from Namani et al. 2019 and Rube et al. 2016 were run through all software combinations in Rocketchip three times each. Raw peak counts were log2 transformed and plotted in the heatmap. Darker red corresponds to higher peak counts while darker blue corresponds to lower peak counts. Gray corresponds to “NA” values, as PePr and CisGenome cannot be run without a control. The heatmap was created using R (v4.2.3) with a kernel (r-irkernel v1.3.2) in Jupyter Notebook (v1.0.0) using the following packages: ComplexHeatmap (v2.14.0), dplyr (v1.1.4), tidyr (v1.3.1), reshape2 (v1.4.4), stringr (v1.5.1), and MASS (v7.3.60.0.1).

For the Namani data, notable contrasts in peak-calling outcomes were observed across various algorithms. Genrich consistently yielded the highest number of peaks, contrasting with the Rube data where MACS3 showed higher peak counts. Interestingly, PePr consistently produced the lowest peak counts for the Namani dataset despite performing second best for the Rube data. The choice of aligner did not significantly impact peak counts overall; however, omitting deduplication generally resulted in slightly higher peak counts across all peak-callers. Unlike the Rube data, where the distinction between narrow- and broad-peak calling showed minimal differences across algorithms, the Namani data exhibited variability based on this distinction. Specifically, MACS3 identified higher peak counts under the assumption of broad peaks compared to narrow peaks.

In addition to assessing the results of the different algorithms, we also assessed Rocketchip’s ability to replicate experimental findings. A total of 288 combinations were tested, accounting for both data sets and software combinations. Among the 288 combinations tested, 274 trials (95.14%) demonstrated perfect replication of peak counts across all three trials. Surprisingly, despite identical software versions, computational resources, and inputs, certain combinations of data and software were non-deterministic, with 4.86% (14 out of 288) exhibiting variability in peak counts (Table 1). The greatest variability in peak counts occurred when CisGenome was used as the peak-caller, where the range between peak counts in these cases were 20,356 and 159,544 peaks. Cases with lower variation (i.e. differences in peak counts ranging from 1-6 peaks) occurred when the STAR aligner was used in the workflow.

**Table 1.**
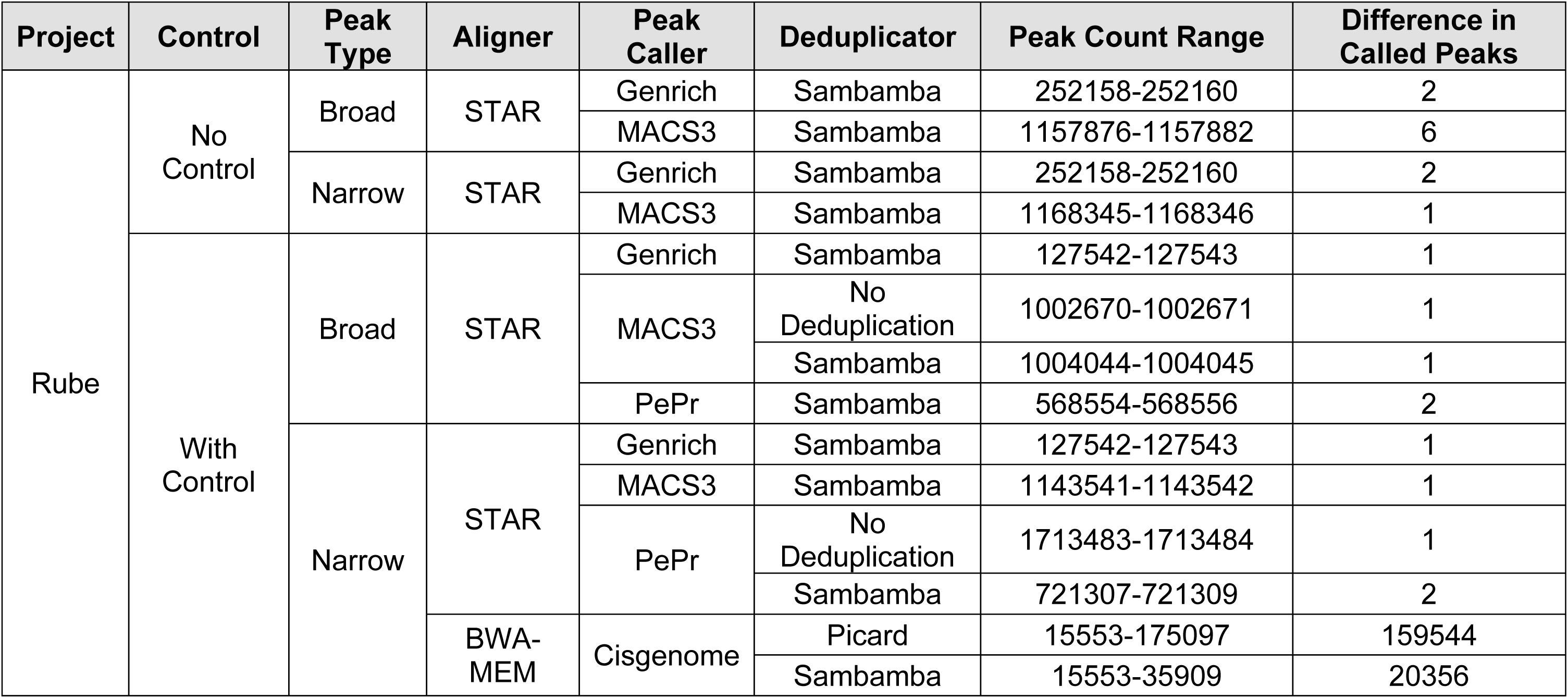
Variation in Called Peaks. ChIP-seq data from Namani et al. 2019 and Rube et al. 2016 were run through all software combinations in Rocketchip three times each. This table depicts all combinations of software and data in which peak counts were not replicated perfectly each of the three runs, exhibiting variation in peak-calling. “Project” details the source of the data set. “Control” refers to whether a control was used or excluded during peak-calling. “Aligner”, “Peak Caller”, and “Deduplicator” correspond to the sequence aligner, peak caller, and deduplicator tool used for the Rocketchip run, respectively. “Peak Count Range” represents the minimum and maximum peak counts for the Rocketchip run. “Difference in Called Peaks” represents the range between the minimum and maximum peak counts, highlighting the magnitude of variation in peak-calling across each of the three trial runs.

Overall, these findings underscore four critical points. First, the choice of software combination, including the algorithm, deduplication method, and aligner, has a significant impact on peak-calling outcomes. Second, even with strict control of factors impacting the analysis, we still observe some variability in peak calling, albeit in a small subset of cases. Third, dataset-specific nuances significantly impact the performance of different software combinations, resulting in a differing consensus on what the “best” software combination is. Finally, the results validate that all available software combinations can be successfully executed within Rocketchip, ensuring flexibility and robustness in ChIP-seq analyses.

### Assessing Run Times for Experimental ChIP-seq Data from Varying Genomes and Read Sizes

The UCSC Genome Browser has seven genomes displayed by default in the “Genomes” tab: human (hg38), mouse (mm10), rat (rn6), zebrafish (danRer11), fruitfly (dm6), worm (ce11), and yeast (sacCer3). Three published ChIP-seq data sets on the SRA with varying read coverage were selected per genome and run through Rocketchip (Figure 3) (39–50). These experiments were run on an HPC with 64 CPUs and 250 GB of memory available. However, jobs were run without being parallelized (i.e. one job at a time with one thread). Additionally, genome copies were deleted between runs using the same genome to ensure that the run time accounts for the full workflow. All data selected was run on Rocketchip for narrow-peak calling using PE data. The software used was BWA-MEM for alignment, Samtools for deduplication, and MACS3 for peak-calling. The results of this experiment validated the use of the seven major genomes from the UCSC Genome Browser and use of sequence data hosted on the SRA. Additionally, it provides users with estimates of how long Rocketchip should run for different genomes to allow for appropriate computational resource requests. Run times ranged from 0.55 hours to 20.54 hours depending on the genome size and read data used.

**Figure 3.**
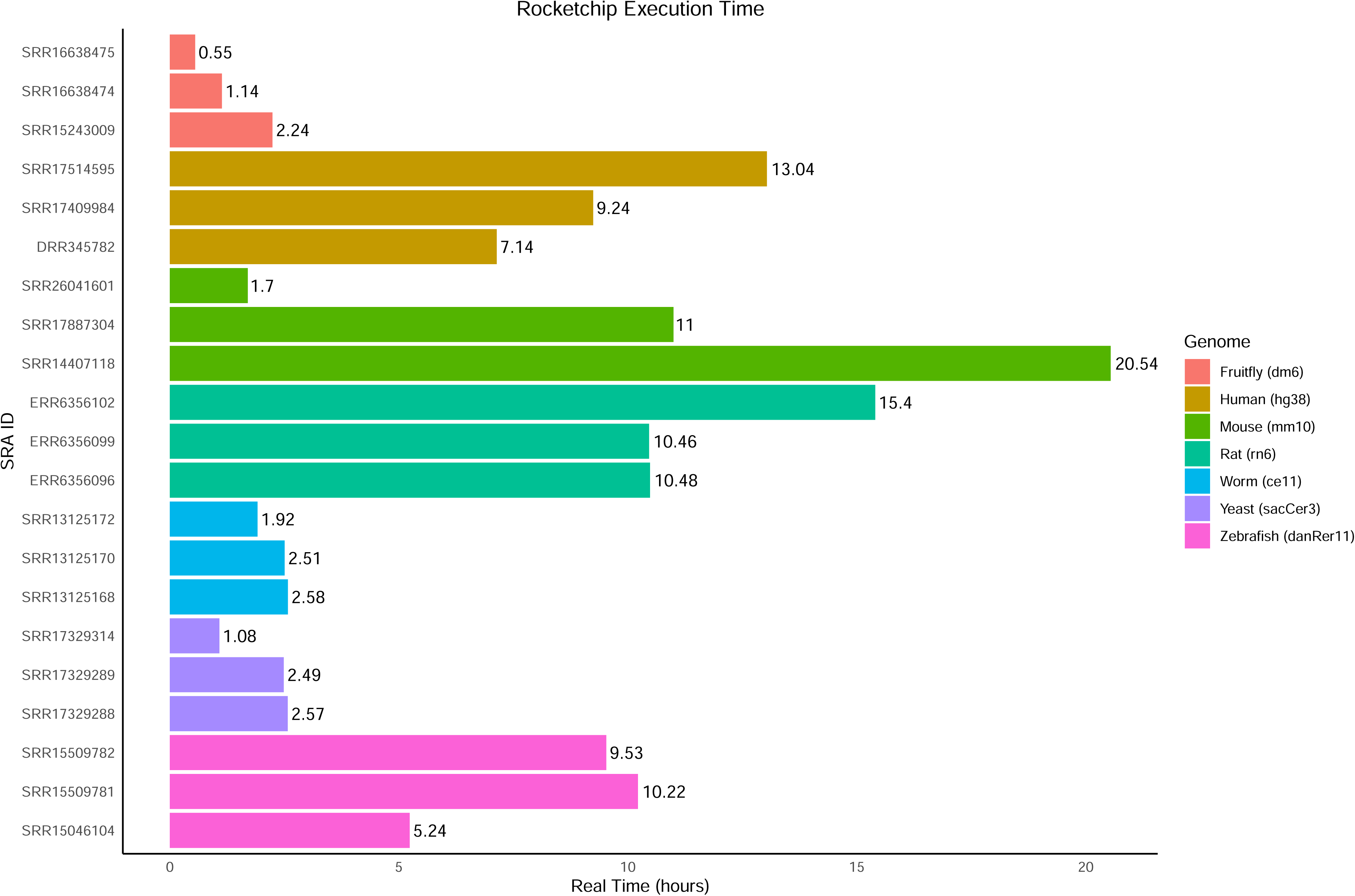
Rocketchip Execution Times per Genome. This is a horizontal bar plot representing how long (in hours) each sample per genome took to run. The label on the bars represents strictly hours (i.e. not hours and minutes). The color of the bars corresponds to which organism the sample comes from and which genome it was run with. The bar plot was created using R (v4.2.3) with a kernel (r-irkernel v1.3.2) in Jupyter Notebook (v1.0.0) using the following packages: ggplot2 (v3.4.4) and dplyr (v1.1.4).

### Validating use of CUT&RUN and CUT&Tag with Rocketchip

As of June 18, 2024, there are 363,213 ChIP-seq, 494,128 CUT&RUN, and 19,492 CUT&Tag data sets available on the SRA. Due to the increasing use of CUT&RUN and CUT&Tag, we wanted to assess Rocketchip’s ability to effectively process data generated by these mapping techniques. Therefore, we applied Rocketchip to CUT&RUN and CUT&Tag data generated by Akdogan-Ozdilek et al. that sought to characterize the zebrafish epigenome during embryogenesis (51). This data set was chosen due to the thorough documentation of the results and high sequence data quality. In evaluating the performance of Rocketchip for CUT&RUN and CUT&Tag data analysis, we assessed the percentage of reads aligned. The alignment percentage was chosen as a metric due to its significance in assessing the overall data processing efficiency and alignment accuracy. Alignment percentage serves as a key indicator of how effectively Rocketchip handles the unique characteristics of CUT&RUN and CUT&Tag data sets, ensuring that a substantial proportion of reads are appropriately mapped to the reference genome and available for further analysis.

The CUT&RUN data set consisted of nine samples. Six samples were SE reads and corresponded to two replicates each for detection of H3K4me3, H3K27me3, and H3K9me3 (SRR14850825 and SRR14850826, SRR14850827 and SRR14850828, and SRR14850829 and SRR14850830, respectively). The remaining three sets were PE read samples that corresponded to two replicates for the detection of RNA polymerase II and mock IP control using the IgG antibody (SRR14850831 and SRR14850832, and SRR14850833, respectively). The study that originally produced these data sets employed Bowtie2 to align sequences, Samtools to filter aligned sequences, and HOMER for peak-calling. Rocketchip was run using Bowtie2, Samtools, and MACS3 for broad peak-calling. As the data was partially SE and partially PE reads, analyses were conducted separately, with the control being used for the PE analysis due to compatibility. Original alignment percentages were compared to those obtained via Rocketchip (Figure 4). A paired t-test was conducted using each SRA input as an observation, yielding a *p*-value of 0.00302, with Rocketchip consistently producing better alignment percentages for the CUT&RUN data compared to the original analysis. The mean difference in the alignment percentages was 32.66%. ‘We hypothesize that the alignment accuracy differences are related to the use of different Bowtie2 versions, as the original study used Bowtie 2.4.1 to align their CUT&RUN data, whereas Rocketchip used version 2.5.2. The GitHub change log for Bowtie2 version 2.5.1, one version earlier than the one Rocketchip uses, notes: “fixed an issue affecting bowtie2 alignment accuracy” and “fixed a segmentation fault that would occur while aligning SRA data.” This highlights the need to revisit and utilize publicly available data, as software updates can improve data processing and thus yield more accurate results. This also highlights Rocketchip’s ability to accurately and effectively process CUT&RUN data.

**Figure 4.**
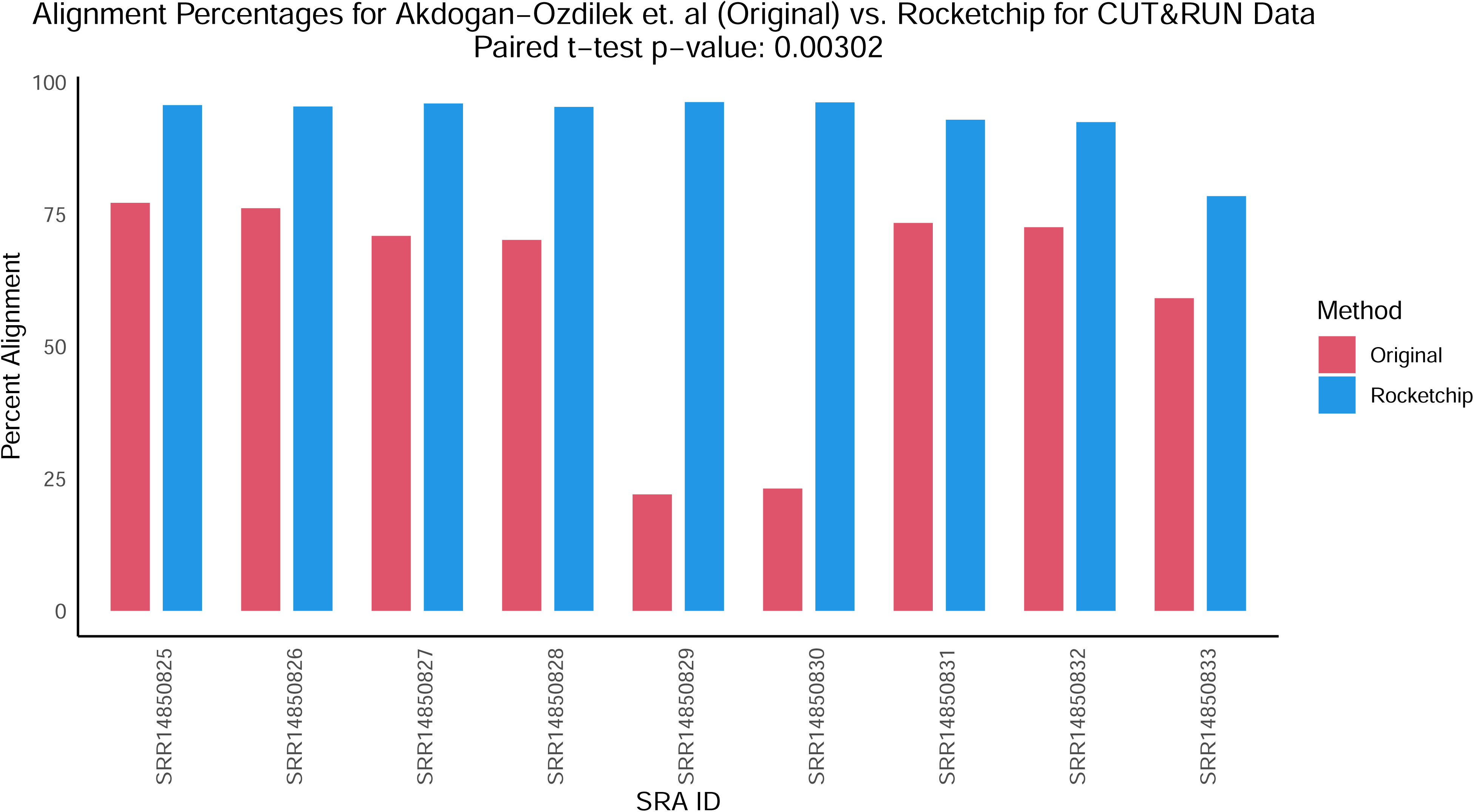
Rocketchip Alignment Percentages for CUT&RUN Data. This is a grouped bar plot comparing the percent of raw reads aligned by Akdogan-Ozilek et al. 2023 (red) compared to Rocketchip (blue) for the CUT&RUN data. The bar plot was created using R (v4.2.3) with a kernel (r-irkernel v1.3.2) in Jupyter Notebook (v1.0.0) using the following packages: ggplot2 (v3.4.4), dplyr (v1.1.4), and tidyr (v1.3.1).

The CUT&Tag data set used for testing in Rocketchip was comprised of six PE read samples. There are three replicates each of H2A.Z at 6 hours and 24 hours post fertilization (SRR14870792, SRR14870793, and SRR14870794 and SRR14870795, SRR14870796, and SRR14870797, respectively). The original study used Bowtie2 for alignment, Samtools for filtering, Picard for deduplication, and MACS2 for peak-calling. We ran Rocketchip using Bowtie2, Samtools, and MACS3. Original alignment percentages were compared to those obtained via Rocketchip (Figure 5). A paired t-test was conducted using each SRA input as an observation, yielding a *p*-value of 0.00015, with Rocketchip resulting in lower alignment percentages for the CUT&Tag data compared to the original analysis. The magnitude of difference in alignment percentages, however, was less than that of the CUT&RUN data, as the mean difference by Rocketchip was -8.41% for CUT&Tag as opposed to +32.66% for CUT&RUN. However, the alignment percentages achieved in our Rocketchip CUT&Tag analysis still surpassed every alignment percentage reported in the original study for their CUT&RUN data, suggesting that Rocketchip is suitable for analyzing CUT&Tag data. We hypothesize that the alignment difference may be due to the usage of both Samtools and Picard in the original CUT&Tag analysis as opposed to just using Samtools in Rocketchip.

**Figure 5.**
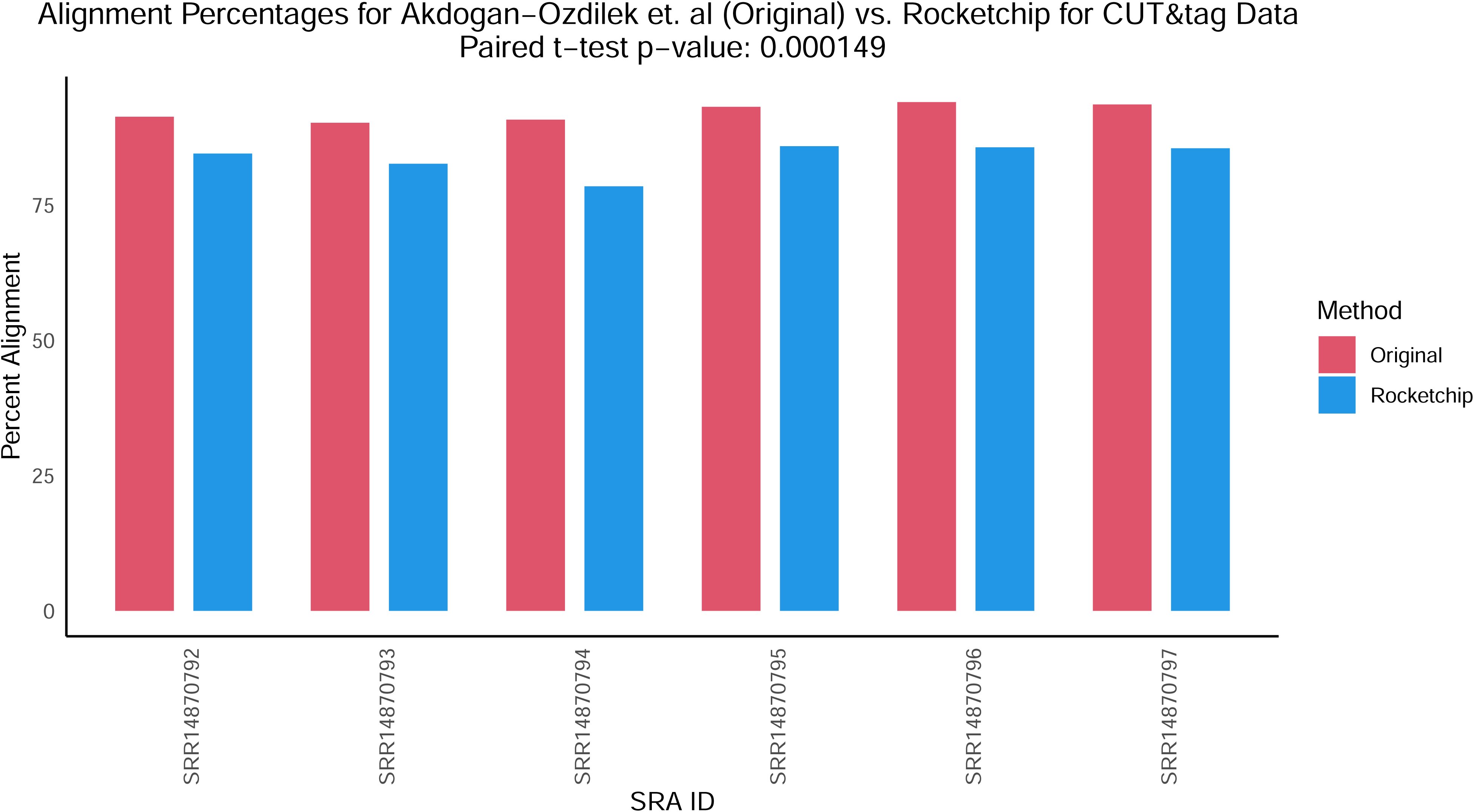
Rocketchip Alignment Percentages for CUT&Tag Data. This is a grouped bar plot comparing the percent of raw reads aligned by Akdogan-Ozilek et al. 2023 (red) compared to Rocketchip (blue) for the CUT&Tag data. The bar plot was created using R (v4.2.3) with a kernel (r-irkernel v1.3.2) in Jupyter Notebook (v1.0.0) using the following packages: ggplot2 (v3.4.4), dplyr (v1.1.4), and tidyr (v1.3.1).

## Discussion

Rocketchip is distinct as a novel and innovative tool due to its unique approach to automating and allowing for flexibility in ChIP-seq, CUT&RUN, and CUT&Tag data analysis workflows. Unlike traditional methods that often require manual intervention and lack reproducibility, Rocketchip provides a straightforward solution by integrating existing software to automatically run analyses for large-scale datasets. Researchers can easily interchange analysis components and rerun their analysis to identify the most appropriate software options for their data. Additionally, Rocketchip was designed to be user-friendly, making it more accessible to researchers with limited bioinformatics expertise compared to traditional methods that require users to navigate software installation, parameter determination, and command inputs from scratch, promoting broader utilization of publicly available sequence data.

In our analyses of Rocketchip using published experimental ChIP-seq, CUT&RUN, and CUT&Tag data, we demonstrated Rocketchip’s ability to handle diverse software combinations seamlessly, which is critical given the variability observed in peak-calling efficiency across different datasets and software tools. We found that the choice of peak caller, aligner, and deduplication method significantly influenced peak-calling outcomes in a data-specific manor. These disparities persisted even when identical software combinations were employed, highlighting the significant impact of dataset-specific factors on the results.

These variations underscore the importance of selecting the appropriate software combination tailored to specific experimental contexts. Moreover, Rocketchip’s ability to replicate results across multiple runs (95.14% perfect replication rate) is noteworthy, demonstrating its reliability in ensuring reproducibility in peak calling. While minor variability (4.86%) was observed in peak counts across some combinations, particularly with CisGenome, this was mitigated with other software tools like MACS3 and Genrich, which exhibited more consistent performance. This is consistent with another study that found that CisGenome performed significantly worse with peak-calling compared to MACS, a prior version of MACS3, as well as having lower consistency in peak-calling (52). Among the other cases where variation was observed in peak-calling, the only consistent variable was that STAR was used as the aligner. There is little documentation to suggest that STAR may yield non-deterministic results, with the exception being a Google Groups thread titled “Reproducibility of alt/ref counts w/STAR alignment (through RSEM)” (53), that suggests that the seed-searching portion of the algorithm is deterministic, but the parallelization (i.e. multithreading) may be the cause of the minor differences observed. Ultimately, the persistence of any variation in results using Rocketchip, which provides controlled software versions and parameters, highlights the need for the further investigation of determinism in algorithms commonly used for analyzing genomic data. Furthermore, there is a seemingly constant push to standardize pipelines and tools, but these analyses demonstrate that standardization is likely not possible due to differences in genomic data. There is, therefore, an increasing need for flexible workflows that provide a streamlined approach to facilitate robust data analysis.

For the CUT&RUN analysis, Rocketchip significantly improved the percentage of mapped reads, likely due to the use of the updated Bowtie2 version. However, for CUT&Tag, Rocketchip resulted in lower read alignment that could not be easily explained by differences in the aligner or deduplicator used prior to peak-calling. This unsolved discrepancy highlights the necessity of documenting software versions and parameters used in analyses to enable replication of results.

When using Rocketchip, a few possible limitations to Rocketchip should be considered. First, it should be noted that broad and narrow-peak-calling must be done in separate Rocketchip runs. This is due to the inherent variability in how peaks are represented via read counts. Similarly, SE and PE data sets must be run separately, as these data types are processed differently at the start of the analysis. Additionally, updating software may yield incompatibilities between dependencies; however, Rocketchip ensures version control via Conda to eliminate potential problems with version incompatibilities.

Future goals for Rocketchip include packaging it to be directly pip or conda installable rather than having a user clone the environment from a YAML file. We are also interested in making a Nextflow adaptation to facilitate cloud computing options. We also look forward to expanding Rocketchip’s selection of software, including other peak-calling algorithms, to provide further user customization options. For instance, WASP (54) has recently become a leading method for deduplication of sequence data. It takes an allele-aware approach to mapping reads back to the reference genome, and discards reads that fail to map to the same region of the genome when the complementary read is considered. This approach reduces false positives for allele imbalance and can help improve peak quality in ChIP-seq data. This algorithm has already been incorporated into STAR alignment. Thus, we aim to release a future version of Rocketchip with an option of using WASP, which will thereby circumvent the following deduplication step in the pipeline. Furthermore, because ChIP-seq can be used for various reasons, such as motif finding or differential binding analysis, automation of further analysis is particularly difficult. For future updates of Rocketchip, we hope to include options for motif analysis via HOMER (55) and differential binding analysis via BEDTools (56) followed by gene ontology enrichments.

Future goals for Rocketchip analyses include using Rocketchip to conduct a meta-analysis of all published data sets for a specific transcription factor. With increased sample sizes and varying coverage, this may yield improved accuracy of transcription factor motifs and *in vivo* binding properties. Additionally, we hope to conduct tests on simulated ChIP-seq data to better understand what factors impact ChIP-seq data and how they do so. This includes modeling narrow vs. broad peak regions, as well as varying peak density and coverage, GC-rich regions, overlapping and bimodal peak regions, and levels of PCR duplication. Synthetic data with these modeled characteristics can be run through the various software combinations within Rocketchip to better understand which tools are better suited for different types of data. Ultimately, this would further researchers’ ability to better tailor specific analysis tools to their data.

## Key Points

- We’ve developed Rocketchip, a Python-based command-line tool that integrates existing software, enabling automated ChIP-seq, CUT&RUN, and CUT&Tag data analysis with ease and efficiency.
- Rocketchip allows for increased leverage of publicly available ChIP-seq, CUT&Tag, and CUT&RUN sequence data.
- Rocketchip is designed to facilitate replicability and reproducibility in molecular analyses of protein-DNA interactions by enabling head-to-head comparisons across published data and software.

## Data Availability

The synthetic sequence data sets, all analysis scripts, and usage instructions can be found in this GitHub repository: https://github.com/vhaghani26/rocketchip_tests. The Rocketchip source code, installation instructions, and usage instructions can be found in the main Rocketchip GitHub repository: https://github.com/vhaghani26/rocketchip.

## Acknowledgements

The authors would like to thank C. Titus Brown, Katie Lee, Brandon Hom, and Aryss Hearne for their computational expertise, statistical expertise, and guidance.

